# ABLE: Attention Based Learning for Enzyme Classification

**DOI:** 10.1101/2020.11.12.380246

**Authors:** Nallapareddy Mohan Vamsi, Rohit Dwivedula

**Affiliations:** BITS Pilani - Hyderabad Campus, India

**Keywords:** attention, bidirectional LSTM, deep learning, enzyme classification

## Abstract

Classifying proteins into their respective enzyme class is an interesting question for researchers for a variety of reasons. The open source Protein Data Bank (PDB) contains more than 1,60,000 structures, with more being added everyday. This paper proposes an attention-based bidirectional-LSTM model (ABLE) trained on oversampled data generated by SMOTE to analyse and classify a protein into one of the six enzyme classes or a negative class using only the primary structure of the protein described as a string by the FASTA sequence as an input. We achieve the highest F1-score of 0.834 using our proposed model on a dataset of proteins from the PDB. We baseline our model against seventeen other machine learning and deep learning models, including CNN, LSTM, BILSTM and GRU. We perform extensive experimentation and statistical testing to corroborate our results.

## 1. Introduction

In any species, enzymes that catalyze reactions in vivo, play a vital role. Their function lies in almost all aspects of biological processes such as nutrition, metabolism and energy conversion. These catalytic agents are grouped together based on the class of reactions they catalyse. It is highly crucial to annotate the enzyme function, because this can help in a wide range of applications such as industrial biotechnology, metagenomics [1], establishment of gene-protein-reaction associations in metabolic pathways [2] and most importantly, determination of enzyme-deficiency related diseases [3]. The currently employed wet laboratory-based experiments for the determination of enzyme class are quite expensive, laborious and time taking. With the number of enzymes registered to the Protein Data Bank [4] increasing at an exponential pace, there is a dire need for the development of in silico tools which would significantly slash the time and cost for such an endeavour. The scope of such a computational tool would be only to give direction and provide a reasonably good estimate of what the enzyme class would be, but the result would definitely have to be validated by conducting experimental work.

Every enzyme is annotated using the Enzyme Commission (EC) system [5, 6] proposed by the International Union of Biochemistry (IUB). The Enzyme Commission system dictates that each enzyme is associated with a number consisting of four parts separated by a period (A.B.C.D), each of which follow a tree structure. Enzyme Commission numbers constitute an ontological system with the purpose of defining, organizing and storing enzyme functions in a curator friendly and machine readable format. The first number (A) represents which one of the six major classes of enzymes the enzyme belongs to, (i) oxidoreductases, (ii) transferases, (iii) hydrolases, (iv) lyases, (v) isomerases, or (vi) ligases. Each of these six major classes diverge into multiple sub-classes which are represented by the second number (B). The third number (C) represents the sub-sub-class of the enzyme and the fourth number (D) represents the third layer classification of the enzyme.

A wide variety of features, such as primary sequence, secondary and tertiary structures, and functional groups were used for this task in the literature [7]. In this study we studied the prediction of enzyme class solely with the help of the primary sequence. Our contributions in this work are as follows:

- A novel **attention-based bidirectional LSTM** model for classifying enzymes into the major classes which outperforms vanilla deep learning and machine learning models was proposed
- Extensive experimentation on the model was conducted through 10-fold cross validation and by running it against a wide range of machine learning and deep learning baselines on a large, publicly available protein dataset
- Usage of SMOTE, to deal with the enzyme class imbalance problem was proposed
- Rigorous statistical testing was performed to validate our results, instead of relying only on differential increases in performance metrics or scores.

## 2. Related Work

A wide range of machine learning methods including nearest neigbour approches (KNN), neural networks, discriminant analysis, and support vector machine have been used for enzyme classification [7].

Tao et al. [8] compared the performace of four algorithms, Linear Discriminant Analysis, Artificial Neural Network, K-Nearest Neighbors, and Support Vector Machine to classify proteins into seven different enzyme classes using features such as Pseudo Amino Acid Composition (PseAAC), Pseudo Position Specific Scoring Matrix (PsePSSM), Topological Indices, Function Domain (FunD), Torsion Angles, Composition, Transition and Distribution (CTD). Whereas Concu [9] developed a multi-task quantitative structure–activity relationship (QSAR) method to classify proteins into the seven major enzyme classes and sub-classes.

Deep learning based methods include the DeepEC which implemented three Convolutional Neural Networks (CNNs) to first classify the protein as an enzyme or not, second for the third-level enzyme commission number and the third for the fourth-level enzyme commission number. In addition to mlDEEPre [11] which uses Position Specific Scoring Matrices (PSSM), One-Hot Encoding and Functional Encodings to conduct multi-functional enzyme classification using a Convolutional Neural Network, Gao et al. [12] designed three parallel deep convolutional neural networks to learn from sequence and structural features to conduct enzyme function prediction and DEEPre [13] which uses only the sequence based features to predict the enzyme class with a deep neural network with convolutional layers following by an LSTM.

Multiple models have been proposed that use traditional learning models such as eCAMI [14], which is a k-mer based tool to classify 390 Carbohydrate-active enzymes (CAZymes) families into thousands of sub-families. Each sub-family with their own distinguishing k-mer peptides. Wheras Dobson and Doig [15] experiments with multiple types of SVM to predict enzyme class. Additionally, Ezypred [16] is a hierarchical classifier with three layers to predict whether a protein is an enzyme or not, its function class and subclass. A variant of KNN, Optimized evidence-theoretic k nearest neighbor (OET-KNN) is used as the core classification model in this model. Furthermore, Amidi et al. [17] explores the usage of nearest neighbour and SVM fusion approaches for the classification of proteins in the PDB. Lastly, ECPred [18] predicted enzyme class using an ensemble of machine learning models in a hierarchy.

## 3. Materials & Methods

In this section we describe the dataset, the preprocessing steps, the classification models, and present an overview of the performance metrics used to compare the effectiveness of various models. Figure 2 presents an overview of the data processing and classification pipeline, while Figure 1 presents an overview of the end to end classification pipeline.

**Figure 1:**
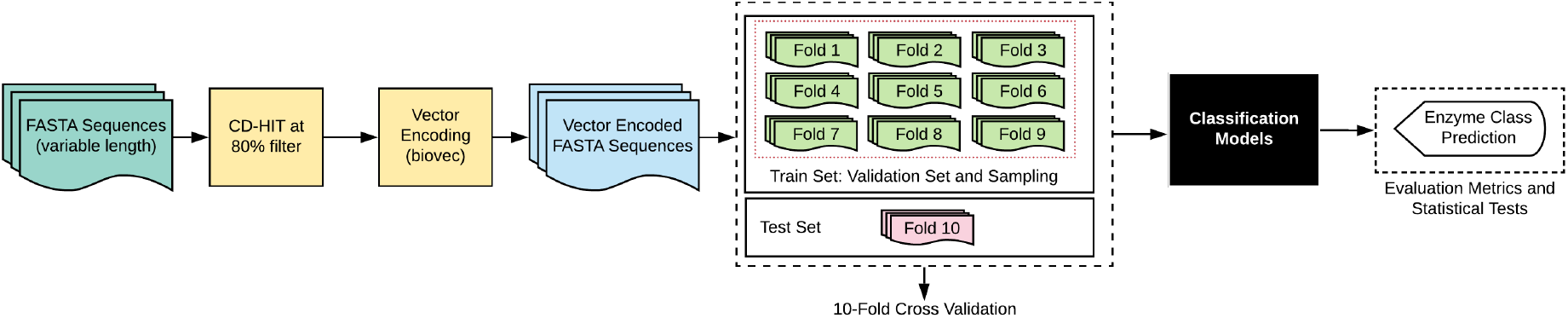
Protein Classification Overview.

**Figure 2:**
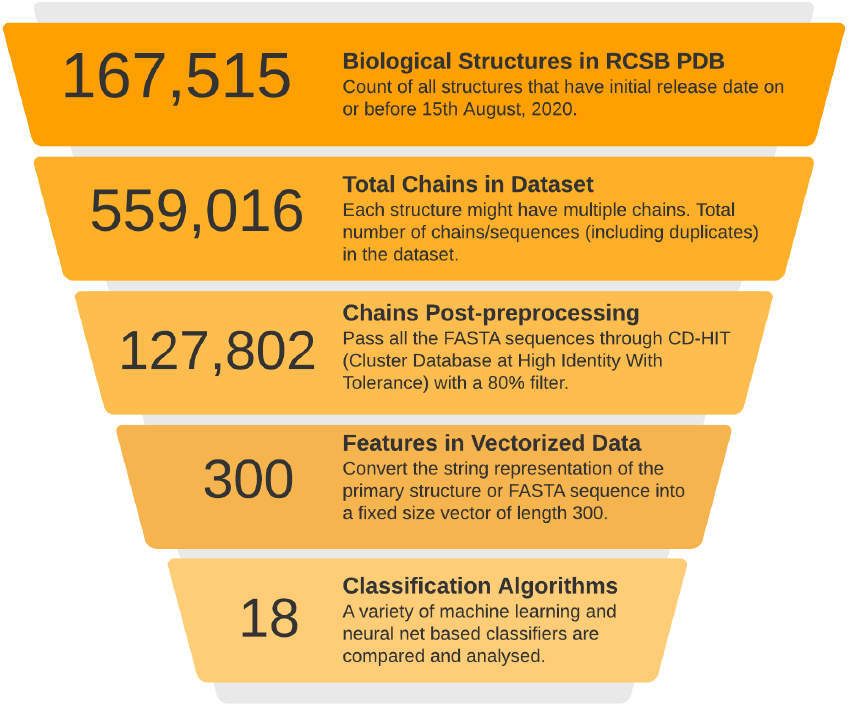
Protein Classification Overview.

### 3.1. Data Compilation and Preprocessing

We use data compiled from the RCSB Protein Data Bank to train, test and benchmark our model^1^. The data is then passed through CD-HIT [19] at a filter level of 30% to remove very similar sequences to reduce bias in the developed models. The dataset consists of 1,27,537 proteins, out of which 91,836 of the them are not enzymes. The remaining 35,701 proteins are enzymes, grouped into one of six classes. As we can see from Table 1, the dataset is rather imbalanced, with the largest class (hydrolases) having six times as many data points as the smallest class (ligases).

**Table 1:** Enzyme Class Frequencies in the Dataset.

| Class | Data Points |
| --- | --- |
| Hydrolases | 10893 |
| Transferases | 10412 |
| Oxidoreductases | 7223 |
| Lyases | 3444 |
| Isomerases | 1880 |
| Ligases | 1776 |

<sup>1</sup>Web link: rcsb.org [4]. We used the RCSB Fetch API to compile all structures in PDB which were released on or before August 15, 2020 (2020-08-15T00:00:00Z).

### 3.2. Data Encoding

Two methods have been used in the related literature to encode the primary FASTA sequence of a protein: (1) one hot encoding and more recently, (2) text embeddings, such as ProtVec or Biovec [20]. One hot encoding of a protein works by identifying that each character in the FASTA sequence of a protein has twenty possibilities - one for each of the twenty amino acids. One hot encoding works by representing each amino acid as a vector of size 20, containing nineteen 0s and a 1 - the position of the one 1 in the vector representing which amino acid it is. For an amino acid with a primary sequence of length *N*, this method would create an encoded output of size 20 × *N*. The second method, which is relatively newer, is based on word embeddings, or Word2Vec [21, 22, 23]. The Word2Vec algorithm trains a shallow neural network with only one hidden layer to learn a lower dimensional projection of the input vector. Once the network is trained, the output layer is removed to retain the connection between the input layer and the hidden layer. This is used to generate projections of the input text. In our method, we used the biovec algorithm which inherently trains a Word2Vec network to generate protein embeddings of the shape 3 × 100.

### 3.3. Cross Validation

We perform a 10-fold cross validation on the dataset with all the baseline models and the proposed models to benchmark and compare the effectiveness of models. 10-fold cross validation works by dividing the dataset randomly into ten segments of equal size. Taking into account the highly class imbalanced nature of the data, we create these folds such that the proportions of each class are same as that of the overall dataset (a *stratified* k-fold cross validation), to ensure that each fold is representative of the complete dataset. In each run, one of the folds (10%)is used as the test set and the remaining nine folds (90%) are used as a train set. For all the deep learning models, we create a validation set using a random 10% of the train set.

### 3.4. Data Sampling

Heavily imbalanced datasets can cause machine learning classifiers to be biased towards the majority class. Sampling methods have been widely used to balance imbalanced datasets, before sending the data to the classification model. Sampling methods can be largely grouped into two categories: (a) oversampling and (b) undersampling. Oversampling methods work by creating new data points of minority classes, while undersampling methods remove data points from the majority classes. One of the most widely used sampling methods is SMOTE, or Synthetic Minority Oversampling TEchnique [24], widely used and customised for various scenarios [25, 26, 27, 28], including biology and health-care use cases like identification of drug-target interaction [29], protein interaction [30] and predicting medical prognosis for ventricular failure [31].

While weighted loss functions have been used to deal with the class imbalance in enzyme classification[15, 12, 13], no other approach has experimented with sampling techniques to the best of our knowledge. We evaluate the performance of all the models: (1) with SMOTE [32], and (2) without SMOTE (no sampling).

### 3.5. Baseline Models

We describe the various machine learning and deep learning models used to baseline and compare our proposed approach. We chose a wide variety of models to be representative of the models that have been used in related literature (Section 2).

#### 3.5.1. Traditional Machine Learning Models

We explore thirteen different machine learning and baseline classification models including AdaBoost [33], Bernoulli Naive Bayes [34], extremely randomized trees [35], k-nearest neighbours (KNN), Gaussian Naive Bayes, Linear Discriminant Analysis (LDA), logistic regression, nearest centroid, passive aggressive classifier [36], perceptron, Quadratic Discriminant Analysis, Ridge Classifer, SGDClassifier, and LightGBM [37, 38].

#### 3.5.2. Deep Learning Models

With the resurgence of interest surrounding neural networks, deep learning [39] has made astounding progress in the last few years. In our study, we developed four different deep learning models which included Convolutional Neural Networks (CNN) [40], Long Short Term Memory Network (LSTM) [41], Bidirectional Long Short Term Memory Network (BiL-STM) [42] and Gated Recurrent Units (GRU) [43]. The parameters for these four models have been mentioned in Table 2.

- **Convolutional Neural Network (CNN)**: A neural network that has one or more convolutional layers which essentially slide a filter over the input. Each convolution filter represents an affine transformation on the input.
- **Long Short Term Memory (LSTM)**: LSTM networks are special recurrent neural network architectures designed to learn long-term dependencies. The additional forget gate allows the network to keep track of information across time steps.
- **Bidirectional Long Short Term Memory (BiL-STM)**: The BiLSTM builds on top of the regular LSTM in the fact that it runs the input in two ways, from past to future and future to past. In this way, the hidden states preserve information both past and future.
- **Gated Recurrent Unit (GRU)**: These are more efficient variants of the recurrent neural network. The advantage comes from solving the vanishing gradient problem using the additional update and reset gates.

**Table 2:** Parameters of the Deep Learning Models.

| Model | Description |
| --- | --- |
| CNN | 3 Convolution Layers [128, 64, 32] and 1 Dense Layer [32] |
| LSTM | 2 hidden layers with 32 nodes each |
| BiLSTM | 2 hidden layers with 32 nodes each |
| GRU | 2 hidden layers with 32 nodes each |

The hidden layers in the CNN model used a Rectified Linear Unit (ReLU) [44] activation function whereas the sequence based networks, LSTM, BiLSTM and GRU, used a Hyperbolic Tangent [45] activation function for the hidden layers. The final output fully connected layer for all the deep learning models used a Softmax activation function. The Adaptive Moment Estimation (Adam) optimizer [46] was used to minimize the Categorical Cross Entropy (CCE) loss function. All the models were trained for 50 epochs with a batch size of 128.

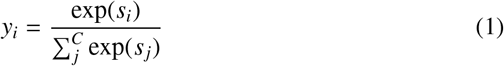

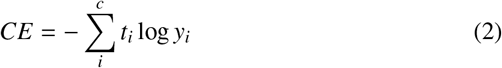

Equation 1 refers to the Softmax activation function. It works in such a way that the summation of the probabilities of all the output classes comes out to be unity. Whereas equation 2 outlines the calculation of the cross entropy (CE) loss.

#### 3.6. Proposed Model: ABLE

We propose an attention based bidirectional-LSTM model used in conjunction with the sampling method SMOTE.

Attention mechanisms (Bahdanau et al. [47]) have been successfully used to enhance performance for a wide range of tasks in sequence data, in natural language processing (modelling sentence pairs [48], relationship classification [49], speech recognition [50]), action recognition [51] and anomaly detection [52]. The attention mechanism relaxes the assumption that each state in a recurrent neural network bears information of the input up until that point. Accordingly, information of all the hidden states corresponding to the whole sequence input is considered and the weights for these hidden states is learned through the following equations. The mechanism has been illustrated in Figure 3, where we can see how the input from each timestep (*h*_*i*_) is assigned a weight (*a*_*i*_) to calculate the context vector (*c*_*t*_).

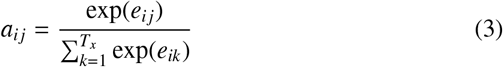

where,

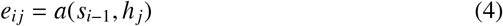

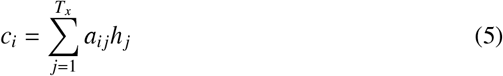

**Figure 3:**
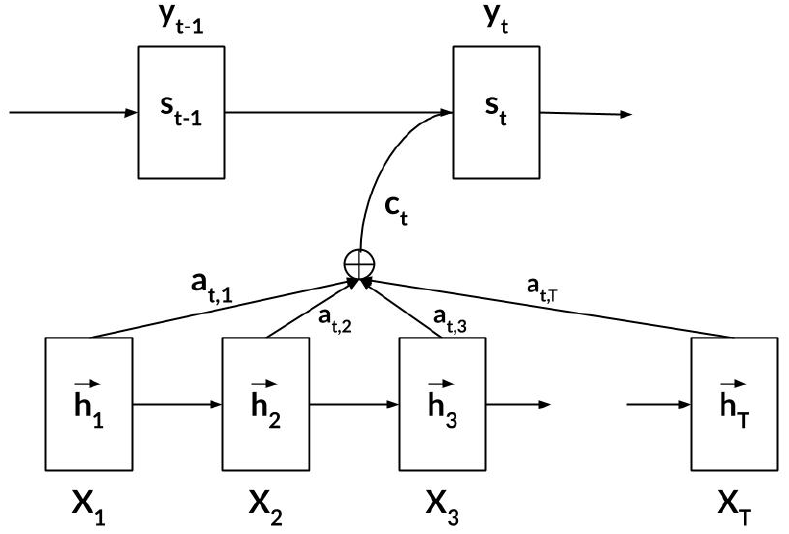
Attention Mechanism Overview.

Equations 3 and 4 showcase how the attention weights are to be calculated and the 5th equation shows how the context vector is calculated. The outlined attention mechanism was exploited to build the ABLE model which is a Bidirectional Long Short Term Memory model with an additional additive attention layer implemented using the Keras Self Attention package [53]. The model has a total of three hidden layers, the first two layers are bidirectional long short term memory layers with 128 nodes which use the Hyperbolic Tangent activation function and the third layer is the Sequence Self Attention layer which also uses a Hyperbolic Tangent activation function. This attention layer is followed by an output fully connected layer with seven nodes, employing a softmax activation function. The model uses an Adam optimizer [46] with a categorical cross entropy loss function.

Training of the model was done for 50 epochs with two callbacks, the first one was a model checkpoint call-back which saved the best performing model in terms of validation accuracy after each epoch and the second callback reduced the learning rate after 10 epochs in which the validation accuracy does not improve. A batch size of 128 was chosen for the training. Additionally, we begin with an initial learning rate of 0.001 and reduce the learning rate as the training progresses, whenever the validation loss plateaus.

All code used for running the models and the results described in this paper have been open sourced under the MIT License^2^.

### 3.7. Performance Metrics

To measure the effectiveness of our model and compare it to machine learning and deep learning baselines, we use the metrics of accuracy, precision, recall and F1 score, all of which are defined in Table 3 in terms of true positives (*TP*), true negatives (*TN*), false negatives (*FN*) and false positives (*FP*).

**Table 3:** Evaluation Metrics.

| Metric | Formula |
| --- | --- |
| Precision | $\frac{TP}{TP+FP}$ |
| Recall | $\frac{TP}{TP+FN}$ |
| F1 Score | $\frac{2TP}{2TP+FP+FN}$ |

The F1-score is the harmonic mean of precision and accuracy. The macro-averaged recall, also known as the balanced accuracy [54], is also used as a performance metric for classification problems on datasets having a class imbalance.

## 4. Experimental Results

### 4.1. Experimental Setup

All experiments were performed on an Intel (R) Core (TM) i5-9300H CPU @ 2.40GHz system with Nvidia GeForce GTX 1650 running on Ubuntu LTS 20.04. Python 3.8.5 with TensorFlow GPU 2.2.0 was used for execution.

### 4.2. Results

Table 4 presents the performance of all models in terms of the average precision, recall and f-score, with and without SMOTE, averaged over the ten folds.

**Table 4:** Comparative Performance of Models: Averages.

| Model | No Sampling |  |  | SMOTE Sampling |  |  |
| --- | --- | --- | --- | --- | --- | --- |
|  | precision | recall | F1 | precision | recall | F1 |
| SGD | 0.349 | 0.219 | 0.190 | 0.259 | 0.307 | 0.190 |
| LGBM | **0.933** | 0.465 | 0.579 | 0.466 | 0.680 | 0.534 |
| LR | 0.268 | 0.194 | 0.203 | 0.235 | 0.344 | 0.218 |
| LDA | 0.374 | 0.221 | 0.240 | 0.253 | 0.374 | 0.257 |
| B-NB | 0.211 | 0.246 | 0.180 | 0.201 | 0.259 | 0.164 |
| NC | 0.186 | 0.219 | 0.158 | 0.188 | 0.220 | 0.159 |
| G-NB | 0.174 | 0.178 | 0.044 | 0.174 | 0.181 | 0.046 |
| AdaBoost | 0.313 | 0.148 | 0.129 | 0.218 | 0.297 | 0.204 |
| Ridge | 0.254 | 0.155 | 0.144 | 0.256 | 0.372 | 0.256 |
| KNN | 0.748 | **0.642** | **0.687** | 0.491 | 0.847 | 0.579 |
| QDA | 0.373 | 0.601 | 0.346 | 0.334 | 0.552 | 0.341 |
| Perceptron | 0.302 | 0.203 | 0.166 | 0.239 | 0.280 | 0.168 |
| PAC | 0.322 | 0.171 | 0.156 | 0.221 | 0.250 | 0.160 |
| GRU | 0.405 | 0.181 | 0.185 | 0.412 | 0.619 | 0.471 |
| CNN | 0.504 | 0.204 | 0.222 | 0.739 | 0.795 | 0.763 |
| LSTM | 0.482 | 0.192 | 0.204 | 0.389 | 0.616 | 0.447 |
| BILSTM | 0.531 | 0.209 | 0.232 | 0.445 | 0.685 | 0.510 |
| **ABLE** | 0.578 | 0.228 | 0.261 | **0.839** | **0.831** | **0.834** |

The proposed model, **ABLE with SMOTE sampling** performs the best overall, achieving the highest recall (0.831) and f-score (0.834). Figure 4 provides a summary of the results of the 10-fold validation for models trained with SMOTE sampling in the form of a boxplot of fscores. We note that for *all* deep learning models (GRU, CNN, LSTM and BILSTM), there is a significant increase in model performance when used in conjunction with SMOTE (average increase in f-score of 0.337). Additionally, attention based BILSTM (ABLE) outperforms the BILSTM on all the performance metrics, suggesting that the usage of attention has helped improve model performance significantly.

**Figure 4:**
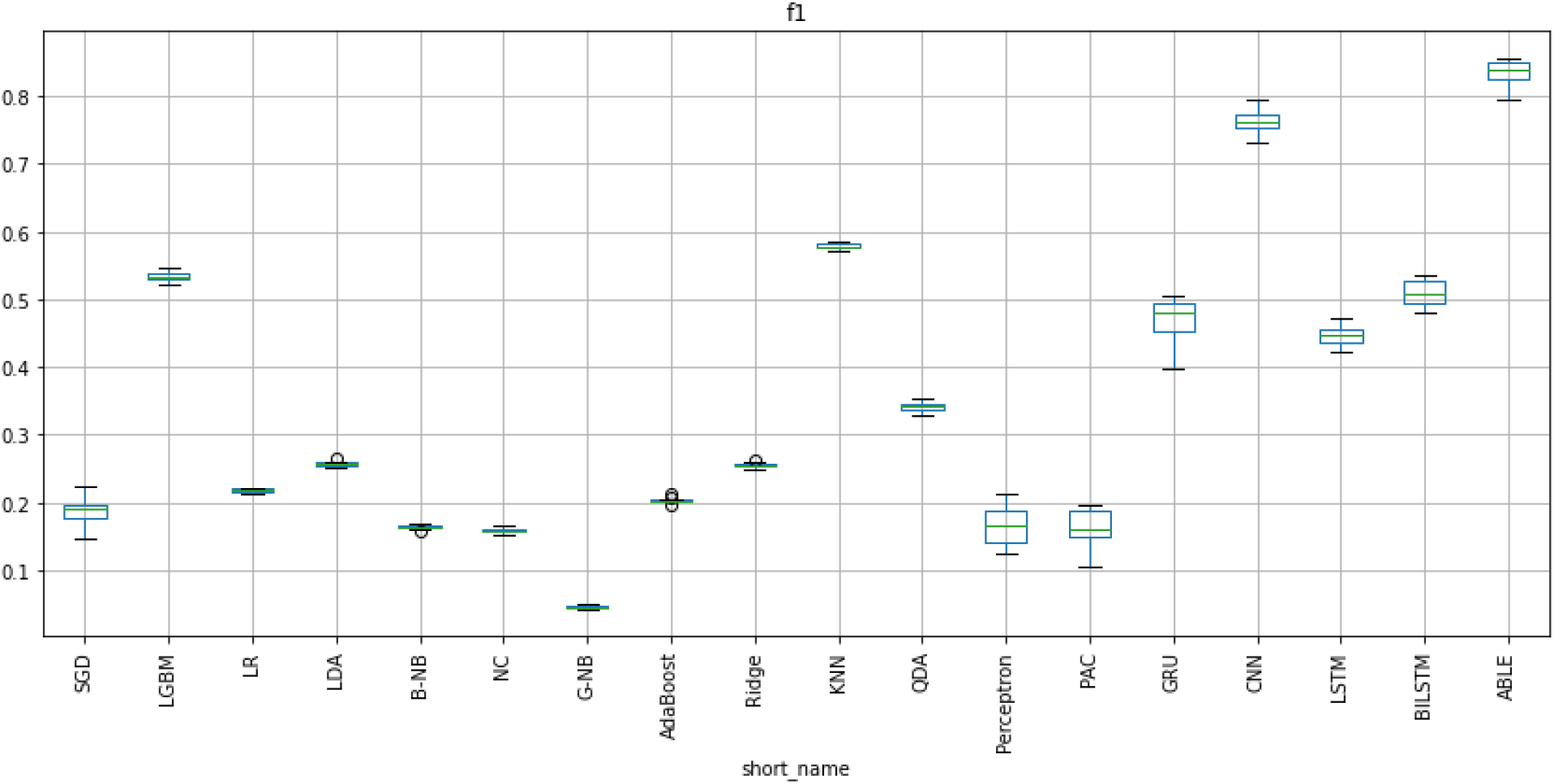
Model performance with SMOTE (f1-scores)

**Figure 5:**
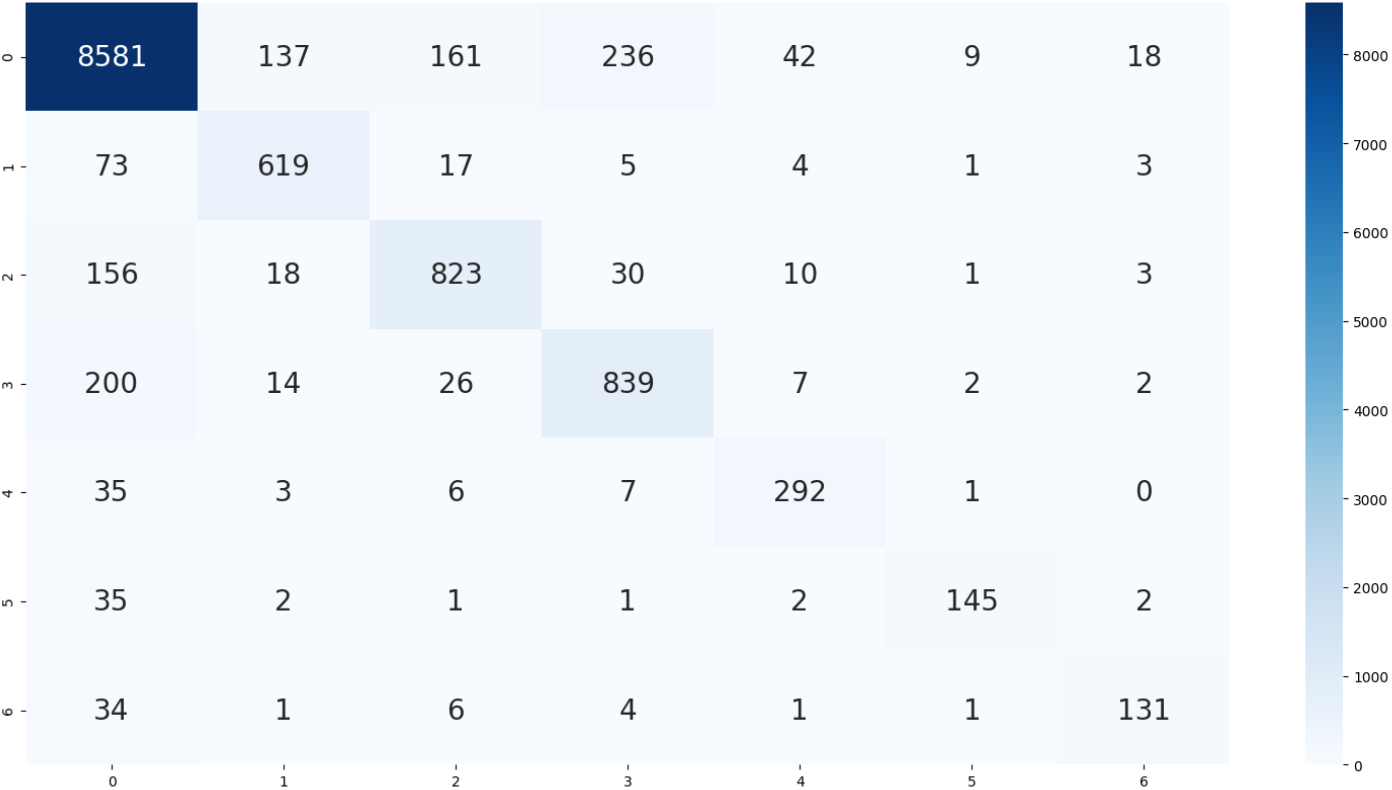
Confusion Matrix on the second fold for the seven classes: Not an enzyme, Hydrolase, Transferase, Oxidoreductase, Lyase, Isomerase, and Ligase.

### 4.3. Runtimes

The summary of execution times of each model, per fold, is presented in Table 5. The values represented correspond to the average time ± the standard deviation. System configuration used for running the experiments has been described in Section 4.1. In addition to the times reported in the table, creating synthetic samples itself using SMOTE was not very computationally expensive and took 41.33 ± 0.46s for each fold (runtime calculated over all ten folds).

**Table 5:** Runtimes of Models (seconds)

| Model | No Sampling | SMOTE |
| --- | --- | --- |
| SGD | $393.876 \pm 4.46$ | $3298.978 \pm 20.31$ |
| LGBM | $415.458 \pm 2.31$ | $1894.660 \pm 3.94$ |
| LR | $174.476 \pm 0.62$ | $816.062 \pm 6.30$ |
| LDA | $73.576 \pm 0.25$ | $399.223 \pm 3.69$ |
| B-NB | $3.606 \pm 0.01$ | $16.968 \pm 0.12$ |
| NC | $0.624 \pm 0.00$ | $3.015 \pm 0.00$ |
| G-NB | $2.397 \pm 0.00$ | $10.834 \pm 0.05$ |
| AdaBoost | $1654.428 \pm 2.95$ | $9803.781 \pm 19.21$ |
| Ridge | $4.745 \pm 0.09$ | $19.007 \pm 0.31$ |
| KNN | $42.600 \pm 0.05$ | $312.665 \pm 1.05$ |
| QDA | $41.591 \pm 0.21$ | $243.032 \pm 4.66$ |
| Perceptron | $34.900 \pm 0.38$ | $182.346 \pm 2.64$ |
| PAC | $56.035 \pm 0.76$ | $265.525 \pm 1.96$ |
| GRU | $1238.13 \pm 15.17$ | $5657.565 \pm 49.70$ |
| CNN | $801.624 \pm 1.04$ | $3888.939 \pm 2.58$ |
| LSTM | $1200.996 \pm 1.91$ | $5563.240 \pm 5.57$ |
| BILSTM | $2004.028 \pm 5.58$ | $9584.289 \pm 52.12$ |
| ABLE | $7578.520 \pm 23.04$ | $37499.214 \pm 88.92$ |

### 4.4. Statistical Test

Existing literature on enzyme classification models compare effectiveness of various models on the basis of raw differences in metrics such as precision, recall or f-score. We perform statistical testing to more robustly understand the comparative effectiveness of the various models used, and to see if the use of SMOTE sampling led to statistically significant improvements in performance.

We use a nonparametric paired test, the Wilcoxin Signed Rank [55, 56] test to analyse the statistical significance of using SMOTE. For comparing the performance of all the models, we perform the Wilcoxon Signed Rank test with *bonferroni correction* pairwise. The *bonferroni correction* is used to solve the multiple comparison problem.

#### 4.4.1. Significance of SMOTE

After dividing the dataset into ten folds, as described in Section 3.3, we run all eighteen models (seventeen baseline models + proposed model) ten times, using a different fold for testing each time. We repeat this procedure with SMOTE and without SMOTE, to form pairs of values, (*m*_*none*_, *m*_*smote*_), where *m*_*none*_ represents the classification metric (precision, recall, or f-score) with no sampling and *m*_*smote*_ represents the performance metric when the same experiment is repeated after using SMOTE on the train data. This totals to 180 pairs of data, on which we perform the Wilcoxin Signed Rank Test. We define the null and alternate hypothesis for this test below:

- **Null hypothesis** (*H*_0_): There is no statistical difference between the two distributions of data observed with SMOTE and without SMOTE.
- **Alternate hypothesis** (*H*_1_): There is a statistical difference between the two distributions of data observed with SMOTE and without SMOTE.

On performing this null hypothesis test with a standard threshold for significance of *p* < 0.05 (significance level, α = 0.05) showed that using **SMOTE is statistically significant** with respect to all three metrics - precision (2.012 × 10^−7^), recall (1.16 × 10^−28^) and f-score (2.55 × 10^−7^), providing more evidence that this improvement in performance is indeed because of SMOTE sampling.

#### 4.4.2. Significance of Models

To compare the performance of models, we perform pairwise tests between all the classification models (*C*_*i*_, *i* ∈ [1, 18]). Since we have 18 models, we need to perform ^18^*C*_2_ = 153 independent null hypothesis tests to compare each pair of models. The null hypothesis in this case will be that for a pair of classifiers (*C*_*i*_, *C*_*j*_), the values of f-scores come from the same distribution, i.e. they are not statistically different. To ensure that we have a significance level of α = 0.05 for each individual hypothesis, we perform a bonferroni correction by rejecting a hypothesis only if the pairwise *p* value satisfies

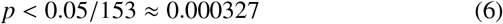

For each null hypothesis test (with models *C*_*i*_, *C*_*j*_, *i* ≠ *j*), we have 20 pairs of observations on which we perform the hypothesis test. Figure 6 summarises the results of the pairwise hypothesis test conducted between the models, with red indicating that the null hypothesis has been rejected, and green^3^ indicating that we have failed to reject the null hypothesis. We note that our proposed model **ABLE is statistically significant with respect to all other deep learning models we baselined against** and is statistically significant with respect to fifteen of the baseline models developed, testifying to the strength of our results.

**Figure 6:**
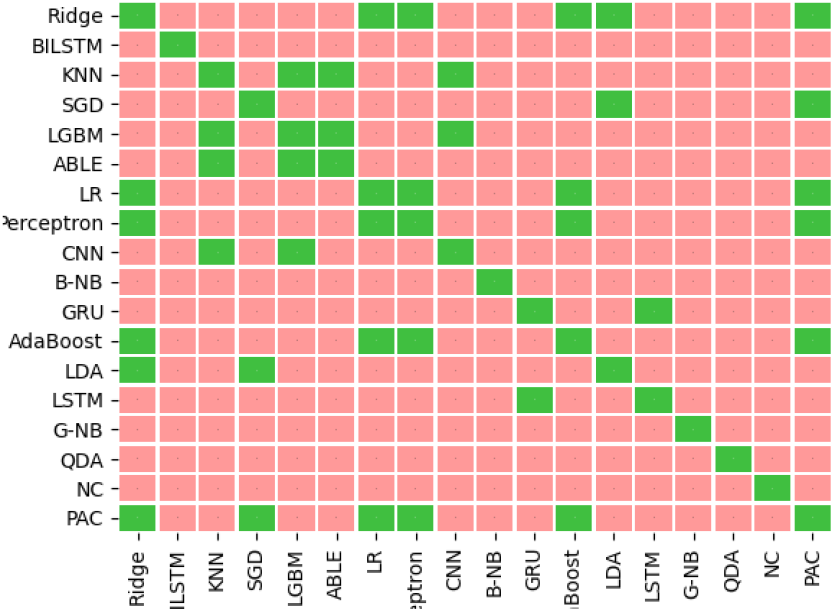
Statistical Significance with respect to the F1 Score.

## 5. Conclusion

We present an attention-based deep learning model for predicting the first part of the enzyme commission number, which uses *only* the primary structure (FASTA sequence) of the protein as an input. We systematically compare attention based models with vanilla deep learning models and machine learning models via 10-fold-validation and statistical testing. It is evident that the usage of deep learning methods incorporating attention can significantly improve the performance of classifiers. Additionally, the use of sampling methods, which has not been seen in literature surrounding enzyme classification before, has shown to be a promising approach to deal with the severe class imbalance problem inherent to enzyme classification systems.

## 6. Funding

This research did not receive any specific grant from funding agencies in the public, commercial, or not-for-profit sectors.

## 7. Declaration of Competing Interest

The authors declare that they have no known competing financial interests or personal relationships that could have appeared to influence the work reported in this paper.

## 8. Author Contributions

**Mohan Vamsi Nallapareddy**: Conceptualization, Software, Writing-Review & Editing; **Rohit Dwivedula**: Methodology, Software, Writing-Original Draft, Visualisation

## Footnotes

1 Web link: rcsb.org [4]. We used the RCSB Fetch API to compile all structures in PDB which were released on or before August 15, 2020 (2020-08-15T00:00:00Z).

2 Link: www.github.com/rohitdwivedula/enzyme-classification

3 All pairs of identical classifiers (*C*_*i*_, *C*_*i*_) have been represented with green in the Figure. However, one must note that the Wilcoxon signed rank test is undefined for identical data, i.e. distributions where the difference between all pairs of data is 0, which is the case for identical classifiers. We use the colour green to indicate there is no difference between the same classifier, and to maintain consistency.

## Notes

https://github.com/rohitdwivedula/enzyme-classification

### Competing Interest Statement

The authors have declared no competing interest.

